# Building an RNA-based Toggle Switch using Inhibitory RNA Aptamers

**DOI:** 10.1101/2021.11.16.468831

**Authors:** Alicia Climent-Catala, Thomas E. Ouldridge, Guy-Bart V. Stan, Wooli Bae

## Abstract

Synthetic RNA systems offer unique advantages such as faster response, increased specificity, and programmability compared to conventional protein-based networks. Here, we demonstrate an in-vitro RNA-based toggle switch using RNA aptamers capable of inhibiting the transcriptional activity of T7 or SP6 RNA polymerases. The activities of both polymerases are monitored simultaneously by using Broccoli and Malachite green light-up aptamer systems. In our toggle switch, a T7 promoter drives the expression of SP6 inhibitory aptamers, and an SP6 promoter expresses T7 inhibitory aptamers. We show that the two distinct states originating from the mutual inhibition of aptamers can be toggled by adding DNA sequences to sequester the RNA inhibitory aptamers. Finally, we assessed our RNA-based toggle switch in cell-like conditions by introducing controlled degradation of RNAs using a mix of RNases. Our results demonstrate that the RNA-based toggle switch could be used as a control element for nucleic acid networks in synthetic biology applications.

**Graphical TOC Entry:** 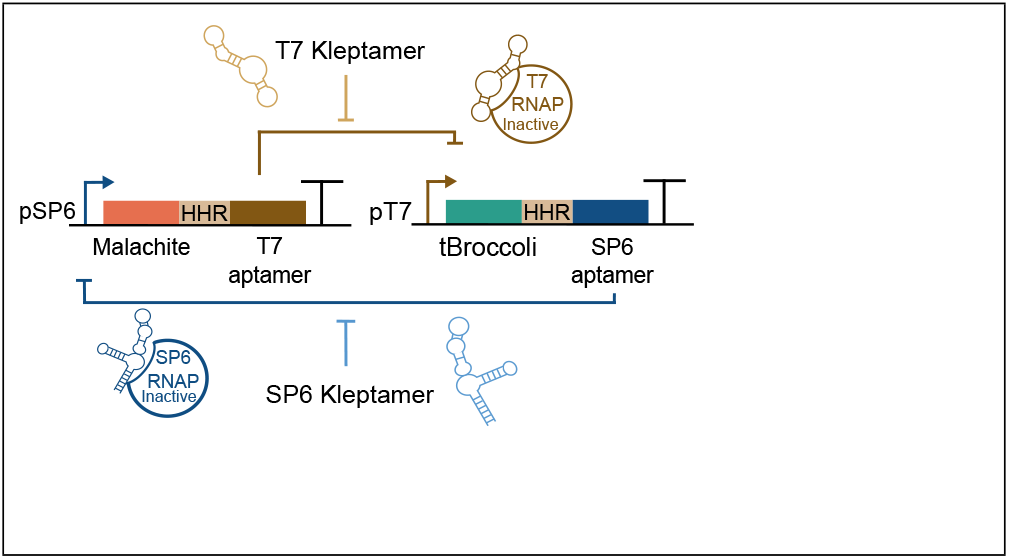

## Introduction

RNA synthetic biology uses RNA-based components to control gene expression and engineer biological systems at the molecular level^1–4^. Compared to their protein-based counterparts, RNA-based circuits impose less cellular burden^5,6^, exhibit faster dynamics, and are typically easier to engineer and tune due to their sequence-specific Watson-Crick base pairing. Consequently, a great deal of attention has recently been devoted to the development of various RNA-based core elements such as riboswitches^7,8^, ribozymes^9^, short interfering sRNAs^10^, RNA aptamers,^11,12^ and riboregulators^13,14^. Among these, RNA aptamers — nucleic acid sequences capable of recognizing and binding to their target molecule with high affinity and specificity — have been used to create RNA-based circuits by interfacing RNA with proteins and other small molecules^15–19^.

RNA-based networks lend themselves to *In vitro* setups. *In vitro* transcription systems have become an attractive alternative to construct and characterize biological circuits without involving the complexities of living systems. With full control over the concentrations and stoichiometries of each component, these platforms are ideal for the rapid prototyping of genetic circuit designs as well as for developing mathematical models to characterize these systems and help fine-tune their design^20,21^.

The most commonly used enzymes for *in vitro* transcription systems are bacteriophage RNA polymerases due to their high processivity for catalyzing the formation of RNA from DNA templates. Among them, T7 and SP6 RNA polymerases (RNAP) are DNA-dependent RNA polymerases that are specific for the T7 and SP6 promoters, respectively. Over the last years, several RNA aptamers have been described to specifically inhibit the activity of the T7 RNA polymerase^22^ and SP6 RNA polymerase^23^. These inhibitory RNA aptamers have been used as regulators for synthetic biological circuits and metabolic engineering applications^24,25^. For instance, Lloyd et al. demonstrated dynamic control of both RNAPs in *in vitro* reactions by applying the principles of strand displacement reactions^26^. They proposed the use of DNA kleptamers - single-stranded DNA oligonucleotides with complementary sequences to the RNA inhibitory aptamers - to characterize minimal circuits with dynamic control over the activation of the RNA polymerases. Here, we take RNA-based circuits to the next level in complexity and bring them closer to a future application in biological systems. We described an *in vitro* RNA-based toggle switch in cell-like degrading conditions. A toggle switch is a bistable network that can be set to either one of two stable states creating a memory unit with an ‘on’ and ‘off’ position. Our circuit is an experimental realisation of a strategy proposed by Mardanlou et al.^27^.

To visualize the transcriptional process in real-time, light-up RNA aptamers are commonly used as a transcriptional reporters. Light-up aptamers are short RNA sequences capable of mimicking the activity of fluorescence proteins. These probes have specific stem-loops that serve as binding pockets for small fluorophores. The quantum yield of the fluorophore significantly increases when it binds to the RNA aptamer^28^. Among them, Spinach and Broccoli aptamers emit green fluorescence signal in the presence of the fluorophore molecule, 3,5-difluoro-4-hydroxybenzylidene imidazolinone (DFHBI)^29–31^. Mango aptamer can bind to thiazole orange molecules to increase fluorophore fluorescence^32^. The Malachite Green aptamer binds to a triphenylmethane dye and emit red (650nm)^33,34^ and the Corn aptamer can dimerize activating the fluorescence of a modified fluorophore from DsRed^35^.To minimise the bleed-through signal between different channels, we used tBroccoli and Malachite Green aptamers.

The architecture of the RNA-based toggle switch here presented is composed of two DNA templates: a first template where a T7 promoter expresses a SP6 inhibitory aptamer and tBroccoli aptamer, and another template where a SP6 promoter produces a T7 inhibitory aptamer and a Malachite Green aptamer (Figure 1). The predicted bistable behaviour of this RNA-based toggle switch arises from the mutual inhibition of transcription by these two RNA aptamers. When an inhibitory RNA aptamer is expressed, it binds to and inhibits the corresponding RNAP, preventing it from binding its cognate promoter. Therefore, two stable states can be reached in this system because when one promoter is active, the other is repressed. These states can be detected by monitoring the fluorescence from the light-up RNA aptamers. In order to reverse the interaction between aptamer and RNAP, specific DNA kleptamers can be added to sequester the inhibitory RNA aptamers and reactivate the RNAPs.

**Figure 1:**
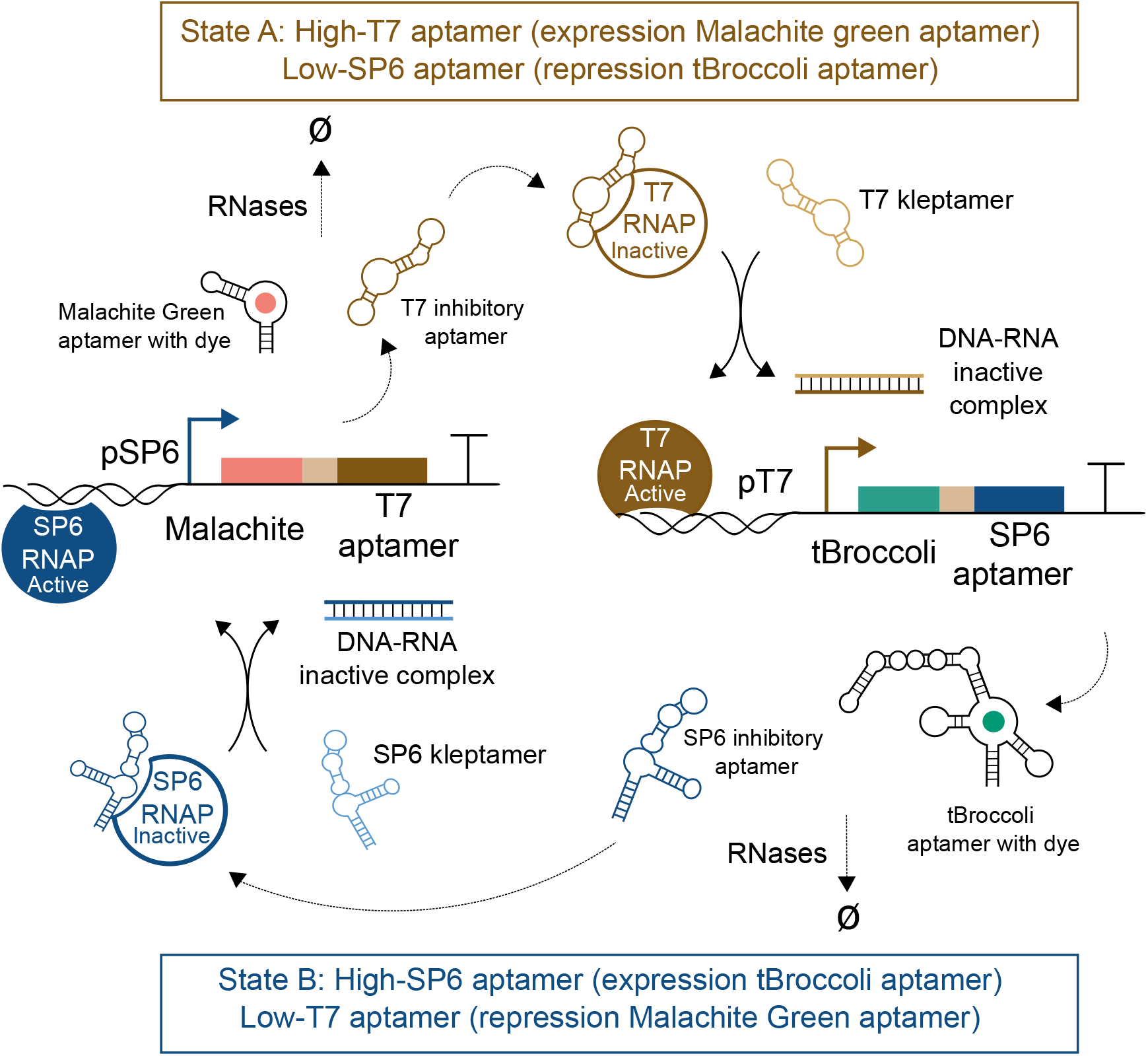
RNA-based toggle switch design. This synthetic network produces two stable states that are reached by the mutual inhibition of the T7 and SP6 inhibitory aptamers. In state A (upper circuit: high-T7 aptamer, low SP6-aptamer), the SP6 promoter drives the expression of Malachite Green and T7 inhibitory aptamer. In this state, Malachite Green and SP6 inhibitory aptamer are repressed. The addition of T7 kleptamers sequester the T7 RNA aptamers and, therefore, reactivate the T7 RNAP that allows the RNA-based circuit to toggle to state B (lower circuit: high-SP6 aptamer, low T7 aptamer). The system can transition back from state B to state A by adding SP6 kleptamers. A hammerhead ribozyme (HHR) is placed between the regulatory aptamer and its associated light-up aptamer (light-brown boxes) to ensure correct RNA folding. RNA molecules are degraded by adding a mix of RNAses.

In this work, we first show that the transcriptional activity of both RNAPs can be suppressed when the inhibitory RNA aptamers are expressed in either cis- and trans-acting circuits. We then demonstrate the recovery of function of RNAPs from cis and trans-acting circuits by adding DNA kleptamers. Next, we show that the RNA-based toggle switch worked at the transcriptional level and can be ‘toggled’ by the addition of DNA kleptamers. Finally, the RNA-based toggle switch proposed is studied in the presence of external RNAses to mimic a cell-like degrading environment for a future application in living cells.

## Results and discussion

### Characterisation of cis-acting RNA-based circuits

We first tested the ability of the inhibitory RNA aptamers to bind to and repress the activity of their RNAP targets. For this purpose, we built two cis-acting circuits: one with the SP6 promoter expressing Malachite Green and SP6 inhibitory aptamers, and another circuit where T7 promoter expresses tBroccoli and T7 inhibitory aptamers. In each DNA template, the inhibitory aptamers are co-expressed along with the fluorescence reporters to monitor the *in vitro* transcription reactions. In the presence of the specific dye, the fluorescence signal of each reporter is detected without significant bleed-through (Malachite Green aptamer - Ex-600 nm/Em-650 nm and tBroccoli aptamer - Ex-472 nm/Em-507 nm). In this work, we decided to separate both molecules by intercalating the hammerhead ribozyme (CChMVd-U10) between the light-up and inhibitory RNA aptamers to improve their folding efficiency and performance^36^.

Both cis-acting circuits showed a significant decrease in the fluorescence signal when the inhibitory aptamers were expressed compared to the expression of only the fluorescent aptamers (Figure 2a,b). The inhibition rates of the SP6 and T7 inhibitor aptamers, indirectly measured by comparing with their associated controls, were roughly 3-fold and 10-fold respectively.

**Figure 2:**
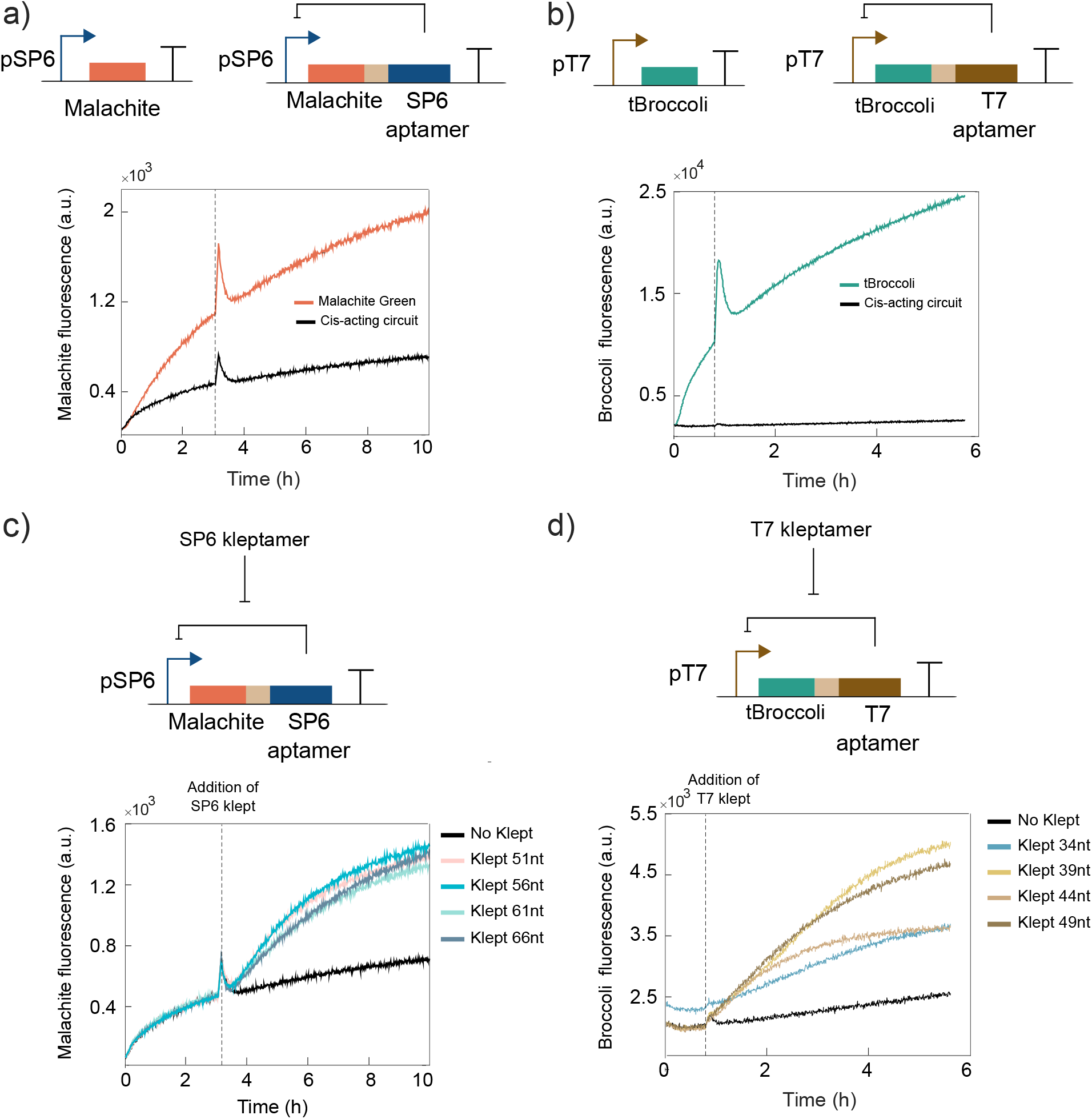
Characterisation of cis-acting RNA-based circuits and DNA kleptamers. **a)** Inhibition analysis of the SP6 inhibitory aptamer in cis-acting SP6 circuits using Malachite Green as reporter. **b)** Inhibition analysis of the T7 inhibitory aptamer in cis-acting T7 circuits using tBroccoli as reporter. **c)** Recovery of Malachite Green fluorescence signal in SP6 cis-acting RNA circuits via addition of 2 *μ*l of SP6 kleptamer.**d)** Recovery of tBroccoli signal in T7 cis-acting RNA circuits via addition of 2 *μ*l of T7 kleptamer. Kleptamer’s injection times are indicated by dashed vertical lines. Spikes in fluorescence are due to injection-induced temperature decrease. Spikes also appear in the inhibition analysis as controls and samples were run at the same time.

After confirming repression of the RNAPs by their cognate inhibitory RNA aptamers, we tested the recovery of transcriptional activity by using DNA kleptamers. We used the Malachite Green aptamer as the reporter for both T7 and SP6 systems and tested the efficiency of DNA kleptamers that covered the sequence of RNA inhibitory aptamers to varying degrees (SI Figure 1). While previous work characterized the DNA kleptamers that cover partially the sequence of the inhibitory aptamers^26^, we designed and tested DNA kleptamers that cover the full sequence of the inhibitory aptamer. Longer kleptamers performed slightly better than the previous reported sequences. We also tested different architectures of the synthetic circuits to observe a significantly better recovery when the fluorescence RNA aptamer was placed 5’ upstream of the ribozyme in the DNA template (SI Figure 1,2,3).

Motivated by these results, we decided to test more designs for DNA kleptamers covering different regions of the sequences. We also incorporate additional binding domains on the kleptamers to test if the dissociation efficiency can be improved by introducing additional binding energy (SI Table 2,3). As shown in Figure 2c, transcription activity of SP6 RNAP was recovered upon addition of 2 *μ*M of DNA kleptamers. Our results indicate that there is no a significant increase in the fold-change between the different designs of kleptamers, but all samples show approximately a 2-fold increase compared to the absence of kleptamers (SI Figure 4).

The activity of T7 RNAP was also recovered after adding 2 *μ*M of the kleptamer (Figure 2d). Although the fluorescence signal increased compared to the control, this signal did not increment as much as would be expected from a free T7 promoter. To test if this effect is due to incomplete release of RNAPs, we tested the efficiency of T7 DNA kleptamers with Malachite Green aptamer (SI Figure 1) to obtain better recovery rates than with tBroccoli reporter. In addition, we observed a similar performance of tBroccoli when it is expressed within the SP6 circuit. We observed a less significant increase with the tBroccoli fluorescence signal using SP6 kleptamer (SI Figure 5). Nonetheless, the recovery rates, normalized with their negative controls were similar for T7 and SP6 kleptamers and different versions of kleptamers showed only a marginal difference in recovering the fluorescence signal. Based on these results, the recovery of SP6 and T7 RNAP’s activity was deemed sufficient for the purpose of constructing an RNA-based toggle switch

### Characterisation of the trans-acting RNA-based circuits

The next step was to assess the performance of both trans-acting circuits that, together, will generate the RNA-based toggle switch. First, we built a circuit in which the T7 promoter expresses tBroccoli and SP6 inhibitory aptamer that, in turn, represses the expression of Malachite Green aptamer in another DNA template (Figure 3a). With the same amount of both DNA templates, we observed repression of Malachite Green aptamer under the control of the SP6 promoter. Only after the addition of the DNA kleptamer, the SP6 RNAP transcribed the Malachite Green aptamer at significant levels while transcription of tBroccoli was not altered (Figure 3b). We also tested that there was no cross-talk between DNA kleptamers by adding consequently both kleptamers (SI Figure 6). We observed that the addition of the incorrect DNA kleptamer had no significant effect in the fluorescence signal of the reporter. This orthogonality is particularly important to ensure the correct performance of the RNA-based toggle switch.

**Figure 3:**
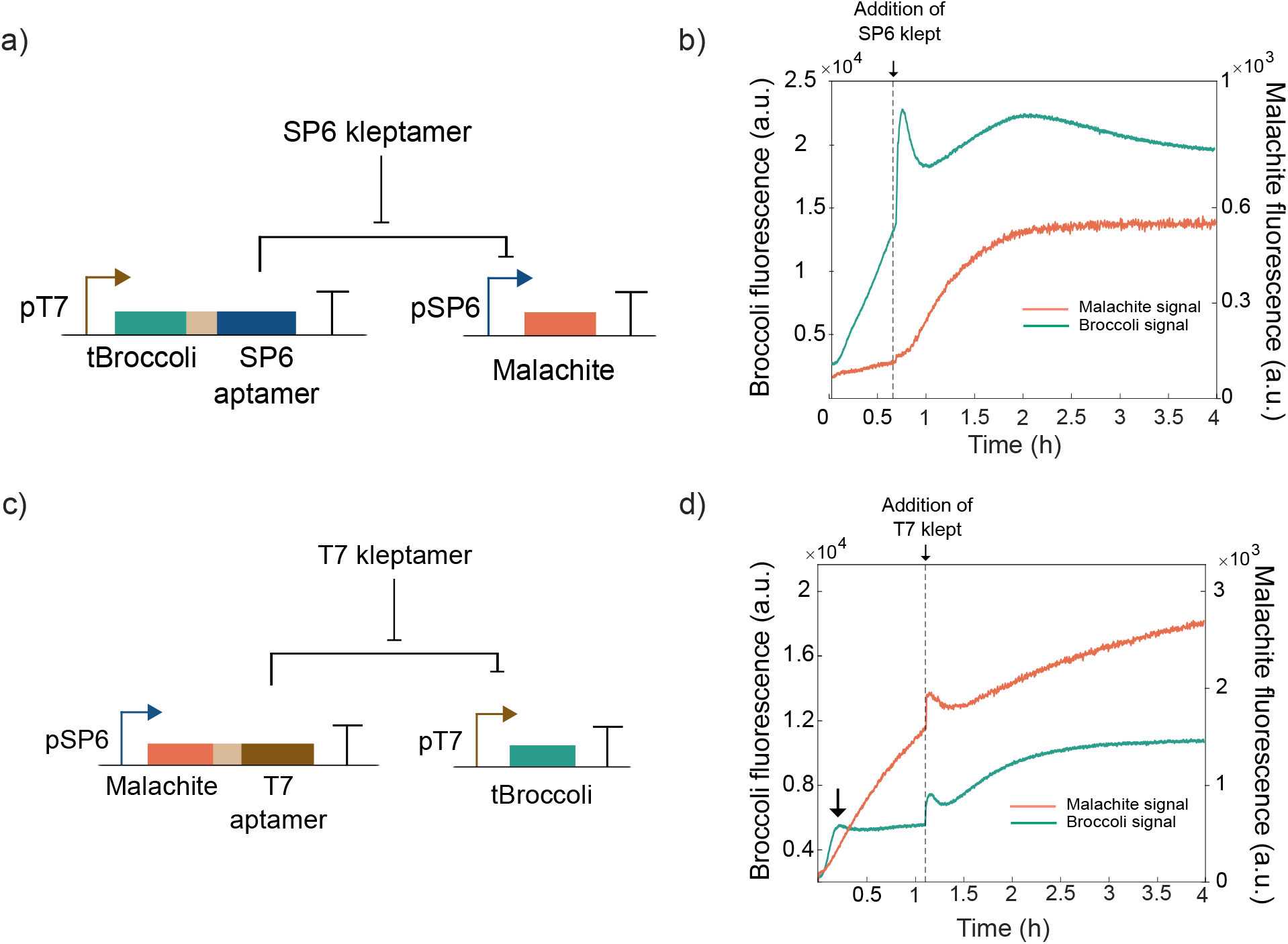
Characterization of individual states of an RNA-based toggle switch. **a)** An SP6 aptamer is transcribed via a T7 promoter, which also produces a tBroccoli aptamer simultaneously. The SP6 aptamer inhibits the expression of a Malachite Green light-up aptamer. **b)** Fluorescence traces demonstrating the inhibition of Malachite green by the system in (a). Only after the addition of SP6 kleptamers (vertical dashed line), Malachite Green fluorescence signal increases. **c)** A T7 aptamer is transcribed via an SP6 promoter, which also expresses Malachite Green aptamer. The T7 aptamer inhibits the expression of a tBroccoli light-up aptamer. **d)** Fluorescence data demonstrates the inhibition of tBroccoli signal. Only after the addition of a T7 kleptamer (vertical dashed line), tBroccoli fluorescent signal increases. Spikes in fluorescent signal near the times indicated by the vertical lines are due to injection-induced temperature changes.

We tested the other trans-acting circuit in which the T7 inhibitory aptamer is produced via an SP6 promoter. T7 inhibitory aptamer represses the expression of tBroccoli aptamer in another DNA template (Figure 3c). When we added the T7 kleptamer to the reaction, the transcriptional activity of T7 RNAP was recovered as expected, as expected by the results of the previous section (Figure 3d). It should be pointed out that the tBroccoli signal increases initially (Figure 3d, black arrow) as the amount of inhibitory RNA aptamer is low at the beginning of the experiment, showing the difference in the strengths of the promoters. The results obtained in this section indicate that a) both T7 and SP6 inhibitory RNA aptamers can be used to repress desired targets in trans-acting circuits and b) trans-acting repression can be mitigated by the use of DNA kleptamers.

### Building a bistable RNA-based toggle switch

We created the RNA-based toggle switch by combining both trans-acting circuits that we characterized (Figure 4a). When we mixed both DNA templates at equal concentrations, we observed domination of transcription from T7 RNAP leading to the repression of SP6 RNAP (Figure 4b, high SP6-aptamer, low T7-aptamer state). This effect is likely due to the difference in the transcription rates between the two promoters. In this state, the system produces tBroccoli and SP6 inhibitory aptamer. The addition of a SP6 kleptamer sequesters the SP6 inhibitory aptamer and frees SP6 RNAP, allowing the expression of the other state of the toggle switch - Malachite Green and T7 inhibitory aptamer -. In this second state, T7 RNAP is repressed and so is tBroccoli signal (low SP6-aptamer, high T7-aptamer state). The subsequent addition of T7 kleptamers sequesters the T7 inhibitory aptamers that are present in the system. This frees the T7 RNAP for expressing tBroccoli and SP6 inhibitory aptamer (high SP6-aptamer and low T7-aptamer state). In this final state, Malachite Green aptamer is repressed by the expression of SP6 inhibitory aptamer while tBroccoli aptamer is expressed. We demonstrate in Fig.4b two subsequent transitions in the same experiment with standard conditions – first from (high SP6-aptamer, low T7-aptamer) state to (low SP6-aptamer, high T7-aptamer) state and back to the (high SP6-aptamer, low T7-aptamer) state.

**Figure 4:**
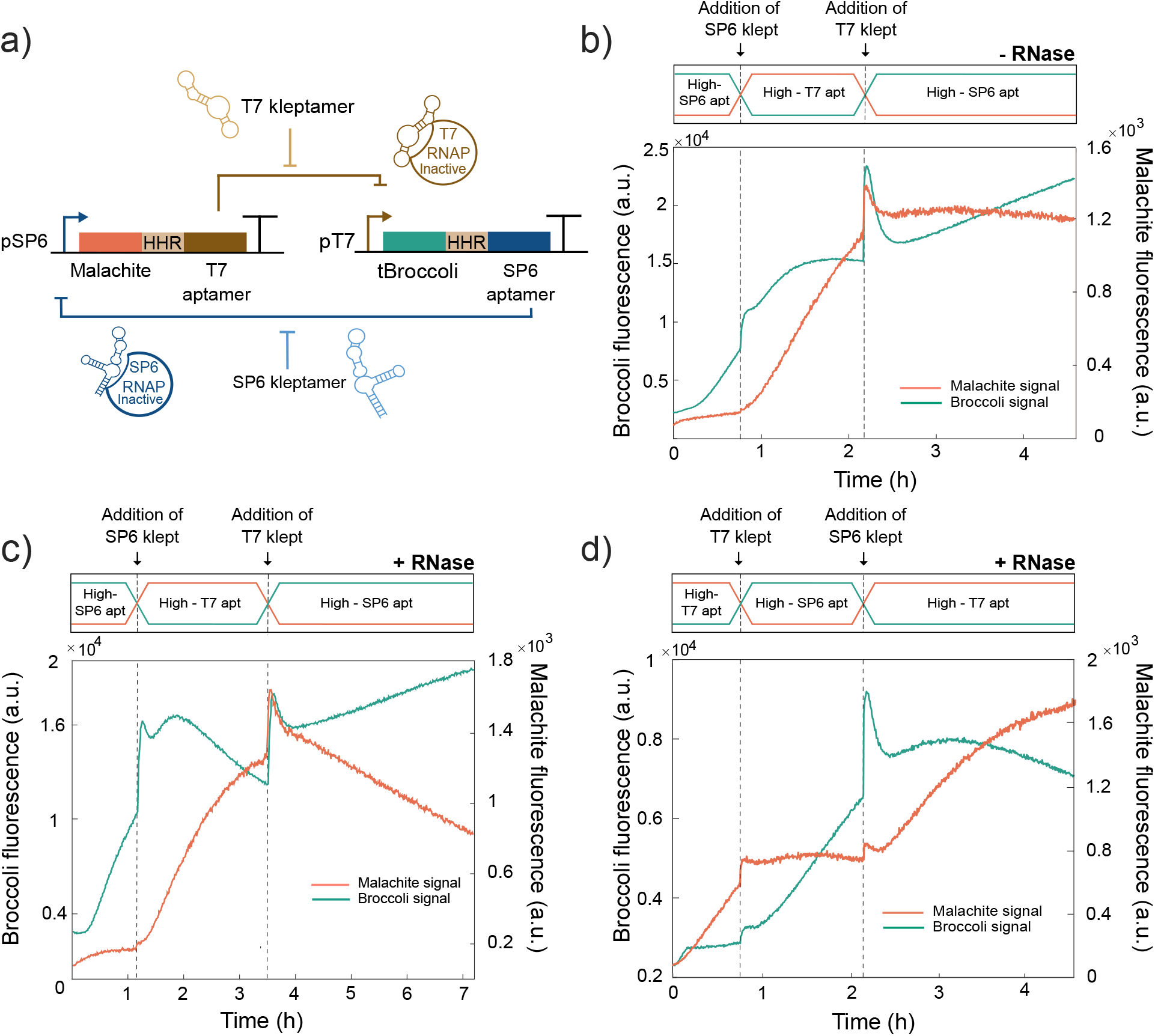
Bistability of RNA-based toggle switch. **a)** Both RNA circuits presented in Figure 3 were combined where T7 inhibitory aptamer is co-expressed with Malachite Green aptamer via a SP6 promoter and SP6 aptamer and tBroccoli via a T7 promoter. **b)** Performance of the toggle switch in standard conditions. tBroccoli fluorescent signal increases initially and after adding the T7 kleptamer whereas Malachite Green fluorescent aptamer only increases after adding the SP6 kleptamer. **c)** Performance of the toggle switch in RNA-degrading conditions. States can be switched after adding the kleptamers and degradation helps pushing to low values the repressed aptamers. **d)** Performance of the toggle switch with an imbalance in DNA templates in RNA-degrading conditions to prove control over this system. The system starts expressing Malachite Green aptamer and can be switched by the addition of the corresponding kleptamers. The vertical dashed lines indicate time points when kleptamers were added. Spikes in fluorescent signal near the times indicated by the vertical lines are due to injection-induced temperature decrease.

Finally, we introduced a mix of purified RNAses in defined amounts to continuously degrade the RNA components in our toggle switch (SI Figure 7). This is an essential feature for a correct operation of the toggle switch and mimicking the live-cell conditions. Also, it pushes down the signal from the suppressed aptamer when its transcription is being repressed. Fig.4c shows the performance of the toggle switch in degrading conditions where the tBroccoli signal is initially increased and then, pushed to low values by degradation in the next state. The degrading environment helps switching between states upon addition of kleptamers by decreasing the concentration of the inhibitory aptamers at repressed states. Finally, as we have full control over the reaction conditions, we introduced an imbalance in the concentration of DNA templates so that the transcription of Malachite Green by SP6 RNAP dominates initially (Figure 4d). We could switch multiple times between states under standard and degradation conditions and with equal or unbalanced concentration of DNA templates.

## Discussion

We have presented the experimental implementation of an *in vitro* RNA-based toggle switch and observed its performance in cell-like degrading conditions. The advantages of RNA molecules including fast dynamics, turnovers, increased specificity and programmability make them ideal for building complex networks for metabolic engineering and synthetic biology. Based on previous works on minimalist RNA-based circuits^26^ and theoretical modeling of an RNA-based toggle switch^27^, we demonstrated the behaviour of this specific network in *in vitro* reactions step by step and highlighted the potential for implementing this network in living cells.

The successful performance of the RNA-based circuits involving RNA inhibitory aptamers and light-up aptamers depends on the secondary structure of the RNA elements. Considering this possible limitation, we decided to separate both RNA aptamers using a self-cleaving hammerhead ribozyme between both structures^37^. We observed better recovery of RNAP activities via DNA kleptamers when the light-up RNA aptamers were placed before the ribozyme (SI Figure 1,2,3). We hypothesize that the residual ribozyme sequence left at 3’-end - self-cleaving occurs near its 5’-end - could have more effects in the secondary structure of light-up aptamers than the inhibitory aptamers. This factor seems to particularly affect the tBroccoli aptamer, where recovery of the fluorescence signal is less significant than for the Malachite reporter after adding the kleptamers. We found that both T7 aptamer and T7 kleptamers performed better when used with Malachite Green aptamer (SI Figure 1). We also found that tBroccoli expressed in the SP6 cis-acting circuit produces even less significant recovery than with the T7 system (SI Figure 5). For these reasons, we believe that RNA secondary structures play an important role in the performance of RNA-based networks *in vitro*. However, this effect could potentially be further improved by the use of other ribozymes^38,39^, catalytic tRNAs^40^ or through engineering the sequence of the HHRs^9^ and also, testing the performance of other RNA fluorescence aptamers such as Spinach^30^, Mango^32^ or Corn^35^. Combining our *in vitro* system with high-throughput measurements would allow the characterization of the system over wider area of operating conditions and parameter values.

To mimic cell-like environment and prove the functionality of the RNA-based system in these conditions, we added a mix of RNases in our system. However, unlike living cells, the rate of *in vitro* transcription slows down as rNTPs are consumed. In order to maintain the activity of RNAPs for longer, we used higher concentrations of rNTPs than standard *in vitro* conditions. With these conditions, RNAPs remain active for 10 hours and we could prove transitions between states by adding kleptamers. However, even 10 hours window was not enough to reach the steady states within each bistable regime in these *in vitro* experiments. By using techniques such as microfluidics, it should be possible to maintain the activity of RNAPs for longer and characterize the behaviour of our circuit more completely. Taking into account the degrading environment of the cell, we used DNA-based kleptamers for building the RNA-based toggle switch rather than RNA-kleptamers. RNA kleptamers could be used for this circuit as well as single-stranded DNA expression systems (SI Figure 8) although more theoretical and experimental work is required to ensure its bistability since, in the presence of RNases, RNA kleptamers would also be susceptible to degradation.

## Methods

### DNA sequences

DNA oligonucleotides and double-stranded DNA (gBlocks™) were synthesized by Integrated DNA Technologies (Coralville, IA) and resuspended in nuclease-free water. Sequences can be found in the Supplementary Information file. NUPACK software was used to computationally check the secondary structures for cis- and trans-circuits. gBlocks were amplified by PCR using Q5 High-Fidelity DNA Polymerase from NEB (NEB #M0492L) and NEB Tm calculator tool was used to estimate the annealing temperature. PCR products were purified using Monarch PCR & DNA Cleanup Kit (#T1030). All Nucleic acid sequences are reported in the SI file.

### *In vitro* reaction mixture

These reactions were performed in 50*μ*l in a transcription mix prepared with 200 ng of each DNA template, 1X RNA polymerase Buffer (NEB #B9012S), 8 mM rNTPs (NEB #N0466S), 250 U T7 RNA Polymerase (NEB #M0251L) and/or 100 U SP6 RNA Polymerase (NEB #M0207L), 15*μ*M of DFHBI-1T and/or Malachite Green dye, and RNase-Free water (Thermo Fisher #AM9932). Malachite Green oxalate salt was purchased from Sigma-Aldrich (Cat. No. M6880-25G) and was dissolved in nuclease-free water and store at −20°C. For tBroccoli aptamer, (Z)-4-(3,5-difluoro-4-hydroxybenzylidene)-2-methyl-1-(2,2,2-trifluoroethyl)-1H-imidazol-5(4H)-one) (DFHBI-1T) was purchased from Bio-Techne Ltd (Cat. 5610/10), resuspended in DMSO and stored at −20°C. RNase Cocktail Enzyme Mix was added with a 1/320.000 dilution factor (Life technologies #AM2286) when relevant.

### *In vitro* transcription experiments

Samples were transferred to a flat *μ*Clear bottom 96-well plates (Greiner). Fluorescent measurements were obtained using a CLARIOstar Plus microplate reader (BMG LABTECH) and reading from the bottom. The temperature of the plate reader was set at 37°C. Fluorescence signals were measured throughout the duration of the experiment. Only at the time of manual addition of kleptamer at a concentration of 2*μ*M, the plate reader was stopped and the plate was outside of the plate reader for less than two minutes. The sample was mixed to ensure the distribution of the DNA kleptamer inside the well. Excitation and emission wavelengths for the dyes were as follows: ex/em= 600/650 nm for Malachite Green light-up aptamer and ex/em= 466.5/512.2 nm for DFHB-1T.

### RNA kleptamers

*In vitro* reactions were performed in 50*μ*l for two hours at 37°C in a transcription mix prepared with 200ng of each DNA template, 1X RNA polymerase Buffer (NEB #B9012S), 8mM rNTPs (NEB #N0466S), 250U T7 RNA Polymerase (NEB #M0251L) and/or 100U SP6 RNA Polymerase (NEB #M0207L), and RNase-Free water (Thermo Fisher #AM9932). GeneJET RNA Cleanup and Concentration Micro Kit (Thermo Fisher #K0841) was used to purify the RNA molecules.

## Acknowledgement

This research is supported by EPSRC grant EP/P02596X/1. T.O. is supported by a Royal Society University Research Fellowship. G.-B.S. is supported by a Royal Academy of Engineering Chair in Emerging Technology for Engineering Biology (CiET1819\5).

## Supporting Information Available

Detailed information on sequences, designs and results.

