## Supplementary material for "Building an RNA-based Toggle Switch using Inhibitory RNA Aptamers": SI Figure

Supplementary Table 1: DNA sequences used in this work.

| **Name** | **Sequence (5'-3')** |
|---|---|
| T7 promoter | TAATACGACTCACTATAG |
| SP6 promoter | ATTTAGGTGACACTATAG |
| tBroccoli | GCCCGGATAGCTCAGTCGGTAGAGCAGCGGAGACGGTC GGGTCCAGATATTCGTATCTGTCGAGTAGAGTGTGGGC TCCGCGGGTCCAGGGTTCAAGTCCCTGTTCGGGC |
| Malachite Green | GGATCCCGACTGGCGAGAGCCAGGTAACGAATGGATCC |
| CChMVd-U10 | GGAAGAGGTCGGCACCTGACGTCGGTGTCCTGA TGATGATCCATGAGAGGATCGAAACCTCTTCTAG |
| T7 aptamer T230-39[a] | AGGCGAGCGTAAGTCAATTCCACTATCATTGCTGCAAGC |
| SP6 aptamer mut9[b] | GGAGAGTTGCTTGGAATGCGTTATAGTCTCTTAGGTGTG TTCGCACACCACTCTCC |

[a] Sequence is taken from Ohuchi et al. [2012] [b] Sequence is taken from Mori et al. [2012]

Supplementary Table 2: DNA kleptamers against T7 inhibitory aptamer

| Name | Sequence (5'-3') |
|---|---|
| T7 kleptamer 32 nt[a] | GAAGCAGCAATGATAGTGGAATTGACTTACGC |
| T7 kleptamer 39 nt | GCTTGCAGCAATGATAGTGGAATTGACTTACGCTCGCCT |
| T7 kleptamer 34 nt | CAGCAATGATAGTGGAATTGACTTACGCTCGCCT |
| T7 kleptamer 44 nt | GCTTGCAGCAATGATAGTGGAATTGACTTACGCTCGCCT **CTAGA**[b] |
| T7 kleptamer 44 nt | **CTTCC**GCTTGCAGCAATGATAGTGGAATTGACTTACGCT CGCCT[b] |
| T7 kleptamer 49 nt | GCTTGCAGCAATGATAGTGGAATTGACTTACGCTCGCCT **CTAGAAGAGG**[b] |

[a] Sequence is taken from Lloyd et al. [2018] [b] Toehold sequences in bold

Supplementary Table 3: DNA kleptamers against SP6 inhibitory aptamer

| Name | Sequence (5'-3') |
|---|---|
| SP6 kleptamer 38 nt[a] | TAAGAGACTATAACGCATTCCAAGCAACTCTCCGCTGC |
| SP6 kleptamer 56 nt | GGAGAGTGGTGTGCGAACACACCTAAGAGACTATAAC GCATTCCAAGCAACTCTCCC |
| SP6 kleptamer 51 nt | GTGGTGTGCGAACACACCTAAGAGACTATAACGCATT CCAAGCAACTCTCC |
| SP6 kleptamer 62 nt | **TCTTCC**GGAGAGTGGTGTGCGAACACACC TAAGAGACTATAACGCATTCCAAGCAACTCTCC[b] |
| SP6 kleptamer 61 nt | GGAGAGTGGTGTGCGAACACACCTAAGAGACTATAAC GCATTCCAAGCAACTCTCC**CTAGA**[b] |
| SP6 kleptamer 66 nt | GGAGAGTGGTGTGCGAACACACCTAAGAGACTATAAC GCATTCCAAGCAACTCTCC**CTAGAAGAGG**[b] |

[a] Sequence is taken from Lloyd et al. [2018] [b] Toehold sequences in bold

Supplementary Table 4: gBlocks

| Name | Sequence (5'-3') |
|---|---|
| T7: T7apt: HHR tBroccoli: T | TAATACGACTCACTATAGAGGCGAGCGTAAGTCAATTCCACTA TCATTGCTGCAAGCGGAAGAGGTCGGCACCTGACGTCGGTGTC CTGATGATGATCCATGAGAGGATCGAAACCTCTTCTAGGCCCG GATAGCTCAGTCGGTAGAGCAGCGGAGACGGTCGGGTCCAGAT ATTCGTATCTGTCGAGTAGAGTGTGGGCTCCGCGGGTCCAGGG TTCAAGTCCCTGTTCGGGCTCACACTGGCTCACCTTCGGGTGG GCCTTTCTGCGTTTATA |
| T7: tBroccoli: HHR: T7apt: T | TAATACGACTCACTATAGGCCCGGATAGCTCAGTCGGTAGAG CAGCGGAGACGGTCGGGTCCAGATATTCGTATCTGTCGAGTA GAGTGTGGGCTCCGCGGGTCCAGGGTTCAAGTCCCTGTTCGG GCGGAAGAGGTCGGCACCTGACGTCGGTGTCCTGATGATGAT CCATGAGAGGATCGAAACCTCTTCTAGAGGCGAGCGTAAGTC AATTCCACTATCATTGCTGCAAGCTCACACTGGCTCACCTTC GGGTGGGCCTTTCTGCGTTTATA |
| T7:tBroccoli | TAATACGACTCACTATAGCCCGGATAGCTCAGTCGGTAGAGC AGCGGAGACGGTCGGGTCCAGATATTCGTATCTGTCGAGTAGA GTGTGGGCTCCGCGGGTCCAGGGTTCAAGTCCCTGTTCGGGCA TGCATGCATGAAAAAAAAAACATGCATGCAGTCGGAAGAGGTC GGCACCTGACGTCGGTGTCCTGATGATGATCCATGAGAGGATC GAAACCTCTTCTAG |
| SP6: SP6apt: HHR : tBroccoli: T | ATTTAGGTGACACTATAGGGAGAGTTGCTTGGAATGCGTTAT AGTCTCTTAGGTGTGTTCGCACACCACTCTCCGGAAGAGGTC GGCACCTGACGTCGGTGTCCTGATGATGATCCATGAGAGGAT CGAAACCTCTTCTAGGCCCGGATAGCTCAGTCGGTAGAGCAG CGGAGACGGTCGGGTCCAGATATTCGTATCTGTCGAGTAGAG TGTGGGCTCCGCGGGTCCAGGGTTCAAGTCCCTGTTCGGGCT CACACTGGCTCACCTTCGGGTGGGCCTTTCTGCGTTTATA |

| Name | Sequence (5'-3') |
|---|---|
| SP6: tBroccoli: HHR:SP6apt: T[a] | ATTTAGGTGACACTATAGGCCCGGATAGCTCAGTCGGTAGA GCAGCGGAGACGGTCGGGTCCAGATATTCGTATCTGTCGAGT AGAGTGTGGGCTCCGCGGGTCCAGGGTTCAAGTCCCTGTTCG GGCGGAAGAGGTCGGCACCTGACGTCGGTGTCCTGATGATGA TCCATGAGAGGATCGAAACCTCTTCTAGGGAGAGTTGCTTGG AATGCGTTATAGTCTCTTAGGTGTGTTCGCACACCACTCTCC TCACACTGGCTCACCTTCGGGTGGGCCTTTCTGCGTTTATA |
| SP6:tBroccoli | ATTTAGGTGACACTATAGGCCCGGATAGCTCAGTCGGTAGAGCA GCGGAACGGTCGGGTCCATCTGAGACGGTCGGGTCCAGATATTC GTATCTGTCGAGTAGAGTGTGGGCTCAGATGTCGAGTAGAGTGT GGGCTCCGCGGGTCCAGGGTTCAAGTCCCTGTTCGGGCGCCAG GAAGAGGTCGGCACCTGACGTCGGTGTCCTGATGATGATCCATG AGAGGATCGAAACCTCTTCTAGACTGGTACGTCCTGCTGATGAG TCCCAAATAGGACGAAACGCGGAAACGCGTCCAGGACTCCACAG TCCGCTCCCATCCTCACACTGGCTCACCTTCGGGTGGGCCTT TCTGCGTTTATA |
| T7: T7apt: HHR Malachite: T | TAATACGACTCACTATAGAGGCGAGCGTAAGTCAATTCCACTAT CATTGCTGCAAGCGGAAGAGGTCGGCACCTGACGTCGGTGTCCT GATGATGATCCATGAGAGGATCGAAACCTCTTCTAGGGATCCCG ACTGGCGAGAGCCAGGTAACGAATGGATCCTCACACTGGCTCAC CTTCGGGTGGGCCTTTCTGCGTTTATA |
| T7: Malachite: HHR: T7apt: T[b] | TAATACGACTCACTATAGGGATCCCGACTGGCGAGAGCCAGGTA ACGAATGGATCCGGAAGAGGTCGGCACCTGACGTCGGTGTCCTG ATGATGATCCATGAGAGGATCGAAACCTCTTCTAGAGGCGAGCG TAAGTCAATTCCACTATCATTGCTGCAAGCTCACACTGGCTCAC CTTCGGGTGGGCCTTTCTGCGTTTATA |

| **Name** | **Sequence (5'-3')** |
|---|---|
| T7: Malachite: T | TAATACGACTCACTATAGGGATCCCGACTGGCGAGAGCCAGGTA ACGAATGGATCCGGAAGAGGTCGGCACCTGACGTCGGTGTCCTG ATGATGATCCATGAGAGGATCGAAACCTCTTCTAGACTGGTACG TCCTGCTGATGAGTCCCAAATAGGACGAAACGCGGAAACGCGTC CAGGACTCCACAGTCCGCTCCCATCCTCACACTGGCTCACCTTC GGGTGGGCCTTTCTGCGTTTATA |
| SP6: SP6apt: HHR Malachite: T | TTTAGGTGACACTATAGGGAGAGTTGCTTGGAATGCGTTATAGT CTCTTAGGTGTGTTCGCACACCACTCTCCGGAAGAGGTCGGCAC CTGACGTCGGTGTCCTGATGATGATCCATGAGAGGATCGAAACC TCTTCTAGGGATCCCGACTGGCGAGAGCCAGGTAACGAATGGAT CCTCACACTGGCTCACCTTCGGGTGGGCCTTTCTGCGTTTATA |
| SP6: Malachite: HHR: SP6apt: T | ATTTAGGTGACACTATAGGGATCCCGACTGGCGAGAGCCAGGT AACGAATGGATCCGGAAGAGGTCGGCACCTGACGTCGGTGTCC TGATGATGATCCATGAGAGGATCGAAACCTCTTCTAGGGAGAG TTGCTTGGAATGCGTTATAGTCTCTTAGGTGTGTTCGCACACC ACTCTCCTCACACTGGCTCACCTTCGGGTGGGCCTTTCTGCGT TTATA |
| SP6: Malachite: T | TAATACGACTCACTATAGGGATCCCGACTGGCGAGAGCCAGGT AACGAATGGATCCGGAAGAGGTCGGCACCTGACGTCGGTGTCC TGATGATGATCCATGAGAGGATCGAAACCTCTTCTAGACTGGT ACGTCCTGCTGATGAGTCCCAAATAGGACGAAACGCGGAAACG CGTCCAGGACTCCACAGTCCGCTCCCATCCTCACACTGGCTCA CCTTCGGGTGGGCCTTTCTGCGTTTATA |

[a] This gBlock was used as a template to change to the T7 promoter via PCR.
[b] This gBlock was used as a template to change to the SP6 promoter via PCR.

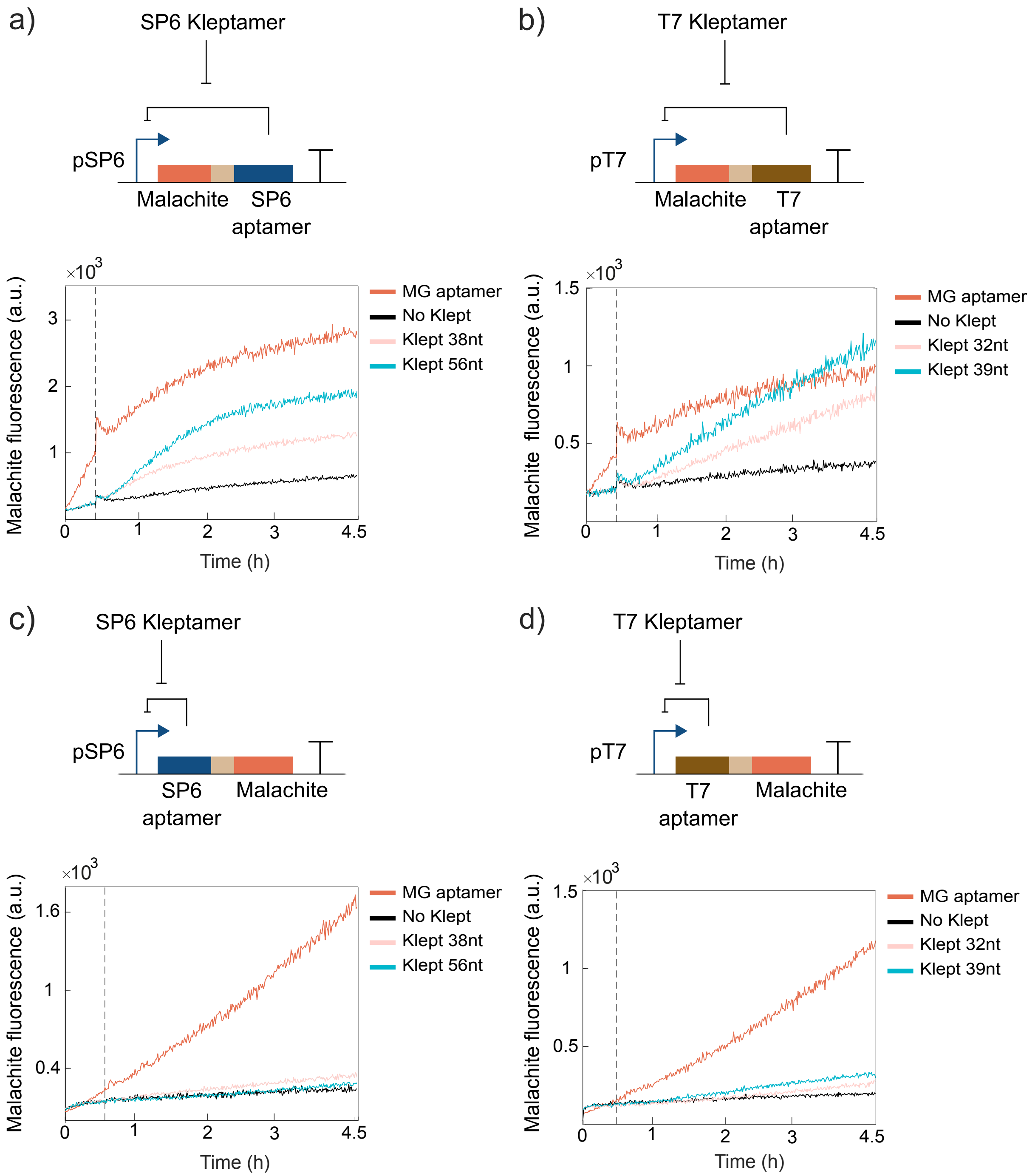


Supplementary Figure 1: Performance of the T7 and SP6 DNA kleptamers in different architectures. DNA kleptamers of 38bp for SP6 RNAP and 32bp for T7 RNAP were obtained from Lloyd et al.. Kleptamers of 56bp and 39bp were designed to cover the full sequence of the inhibitory aptamer. a) SP6 cis-acting circuit and b) T7 cis-acting circuit with the fluorescence aptamer being expressed before the inhibitory aptamer. c) SP6 cis-acting circuit and d) T7 cis-acting circuit with the inhibitory aptamer being expressed before the fluorescence aptamer. Spikes in fluorescence near the times indicated by the vertical lines are due to injection-induced temperature decrease.

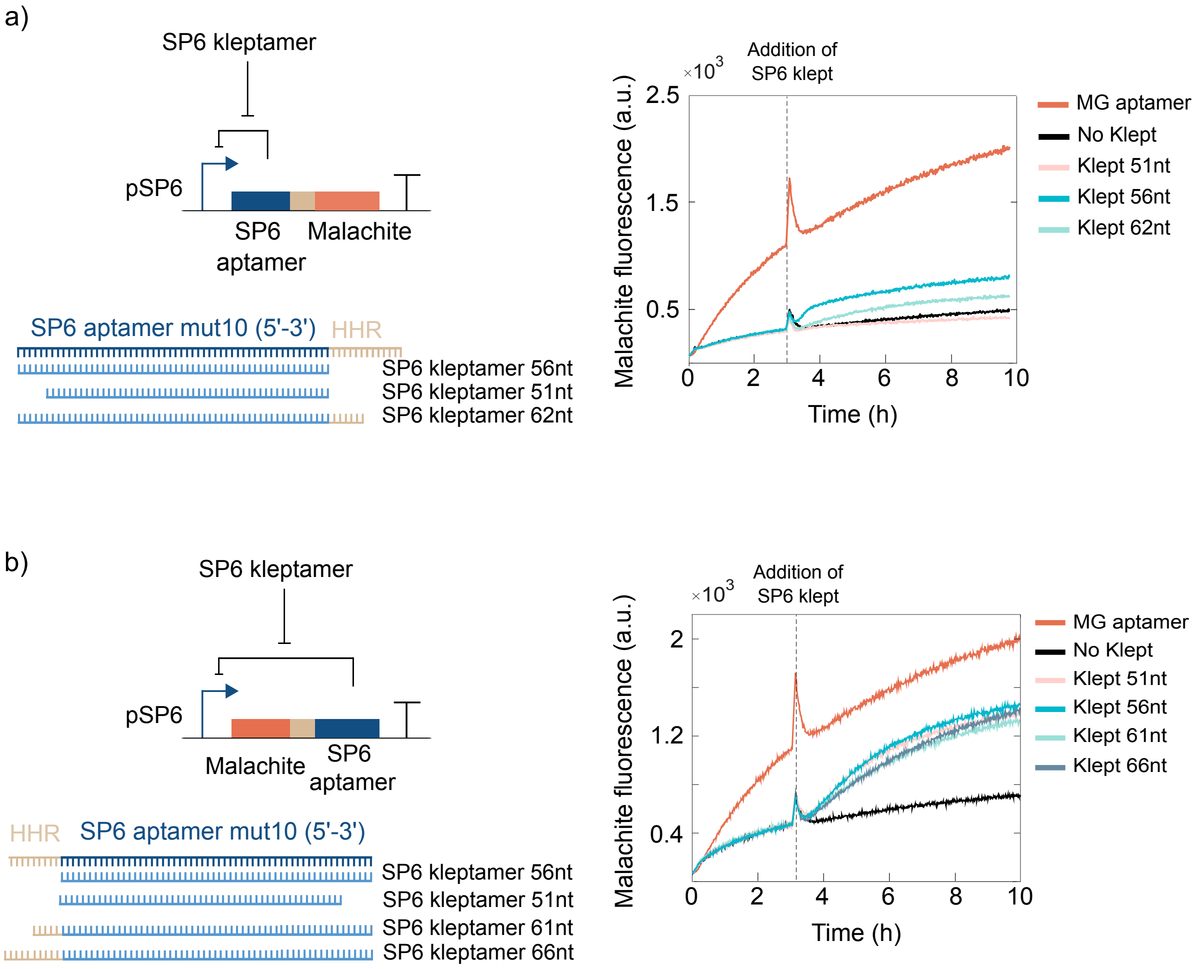


Supplementary Figure 2: Comparison of the performance of SP6 cis-acting circuits with different architectures with their corresponding kleptamers. **a)** Cis-acting circuits where the transcription of the SP6 inhibitory aptamer is followed by the Malachite fluorescent aptamer via SP6 promoter. SP6 kleptamers covering the full and partial inhibitory aptamer were used as well as with a short toehold downstream. **b)** SP6 cis-acting circuits where the transcription of the Malachite Green aptamer is followed by the SP6 inhibitory aptamer. Red lines indicates the expression of only Malachite Green fluorescence signal showing the cost of expressing more elements in the same transcript. Spikes in fluorescence near the times indicated by the vertical lines are due to injection-induced temperature decrease.

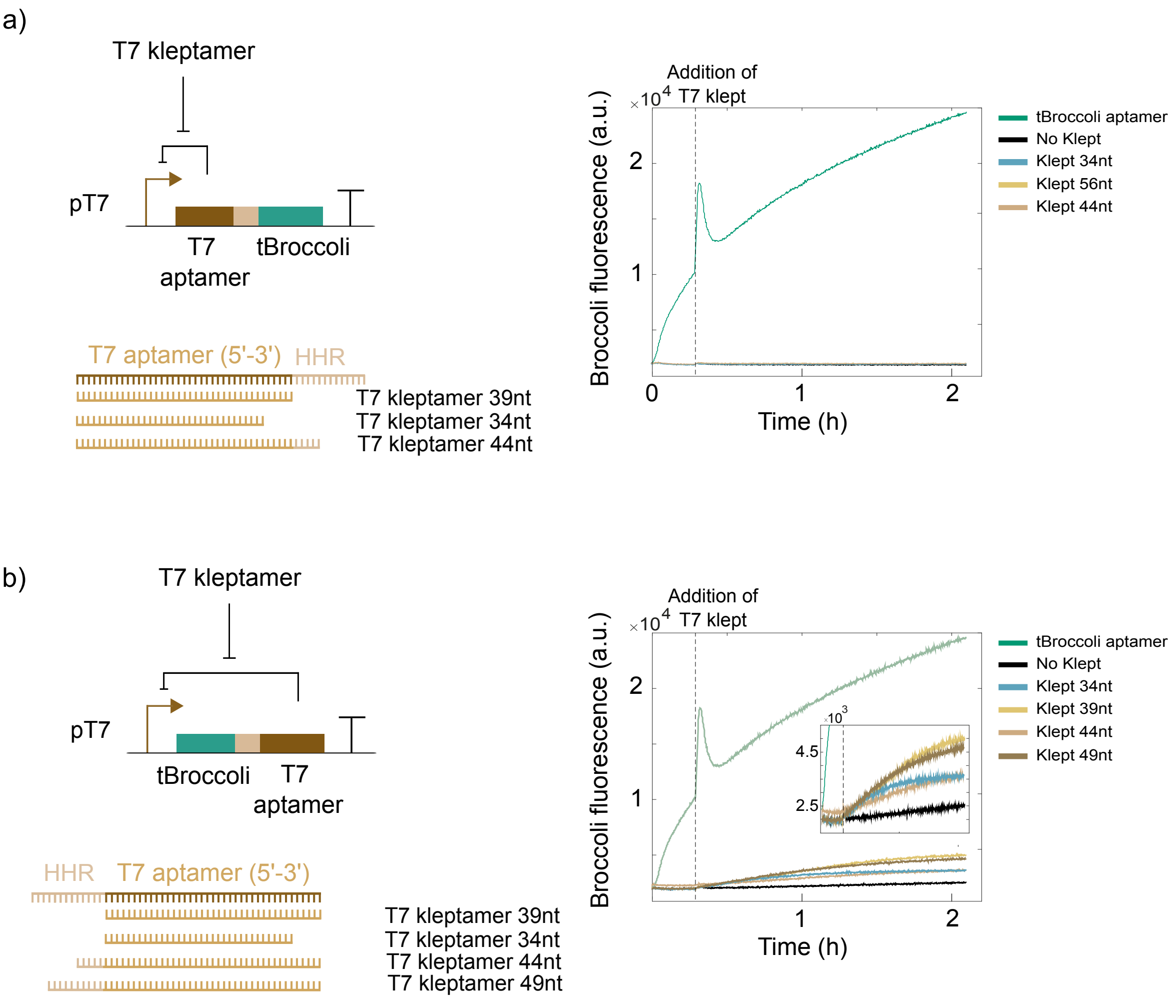


Supplementary Figure 3: Comparison of the performance of T7 cis-acting circuits with different architectures with their corresponding kleptamers. **a)** Cis-acting circuits where the transcription of the T7 inhibitory aptamer is followed by the tBroccoli fluorescent aptamer via T7 promoter. T7 kleptamers covering the full and partial inhibitory aptamer were used as well as with a short toehold downstream. **b)** T7 cis-acting circuits where the transcription of the tBroccoli aptamer is followed by the SP6 inhibitory aptamer. Green lines indicates the expression of only Malachite Green fluorescence signal showing the cost of expressing more elements in the same transcript. Spikes in fluorescence near the times indicated by the vertical lines are due to injection-induced temperature decrease.

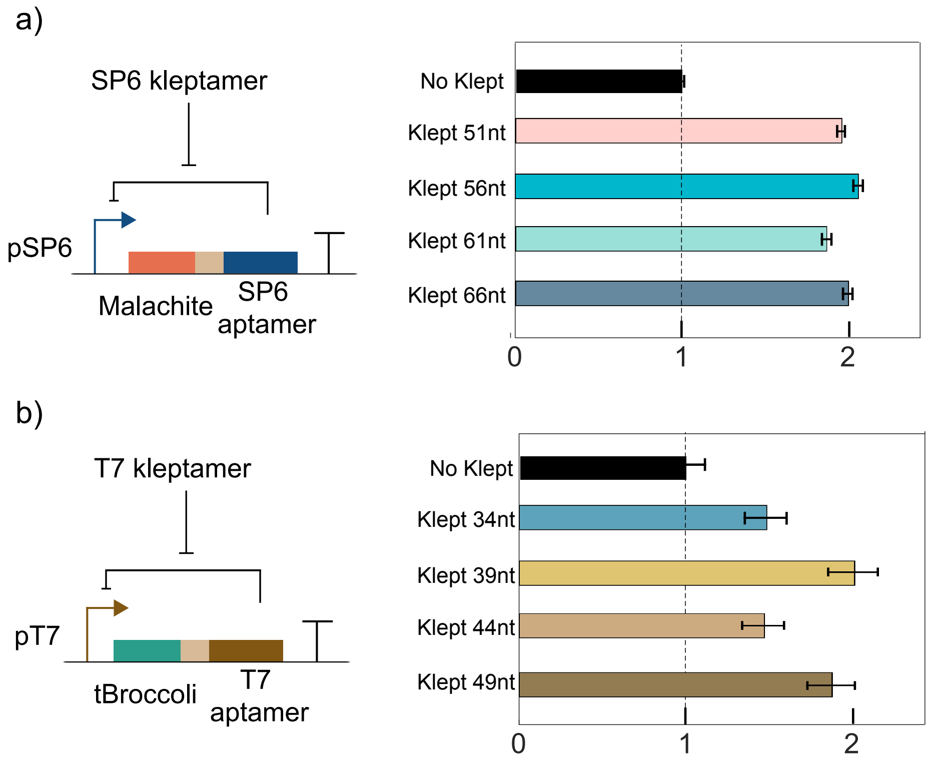


Supplementary Figure 4: Normalization signal from the endpoints of each sample compared to the negative control for **a)** SP6 cis-acting circuits and **b)** T7 cis-acting circuit. The error bars were calculated following the propagation error formula.

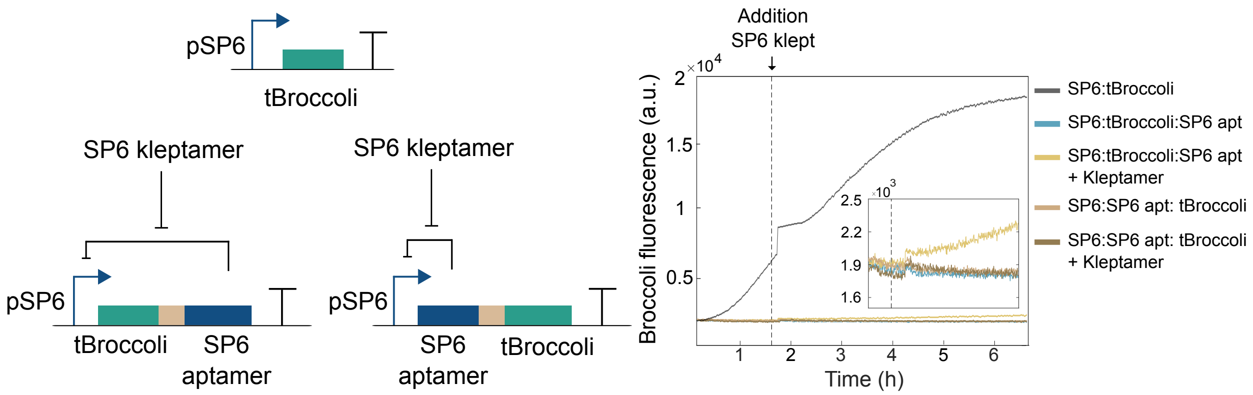


Supplementary Figure 5: SP6 Cis-acting circuit with tBroccoli as reporter. SP6 promoter transcribes tBroccoli and tBroccoli along with the SP6 inhibitory aptamer in both positions. Only when tBroccoli was expressed before the inhibitory aptamer, it was possible to observe an increase in tBroccoli signal after adding SP6 kleptamer. The spike in fluorescence near the times indicated by the vertical lines are due to injection-induced temperature decrease.

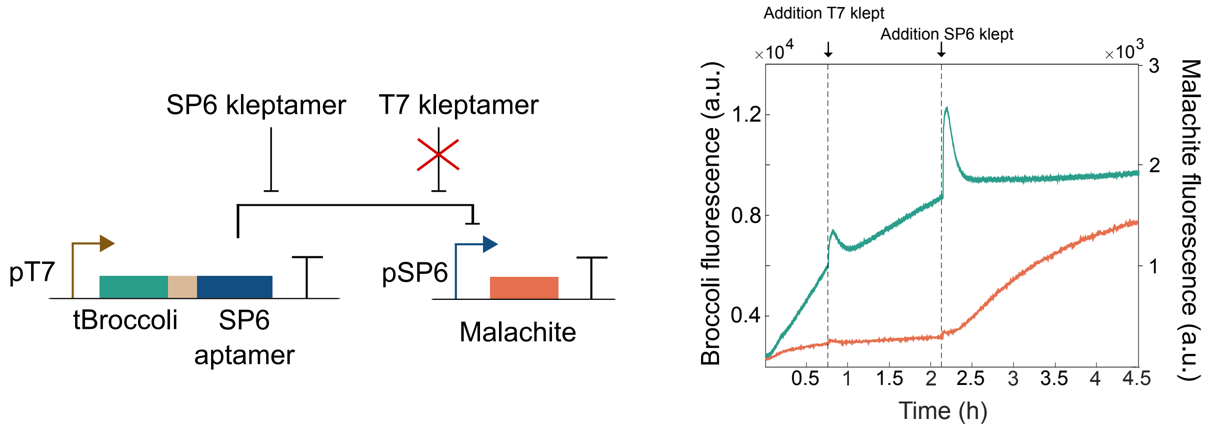


Supplementary Figure 6: Cross-talk between DNA kleptamers tested with SP6 trans-acting circuit. The expression of SP6 inhibitory aptamer represses the production of Malachite Green aptamer. Malachite Green aptamer was still repressed after adding the DNA kleptamer against the T7 inhibitory aptamer (first black vertical dashed line). Only after the addition of the adequate DNA kleptamer, the SP6 RNAP was able to start the transcription of this fluorescent aptamer.

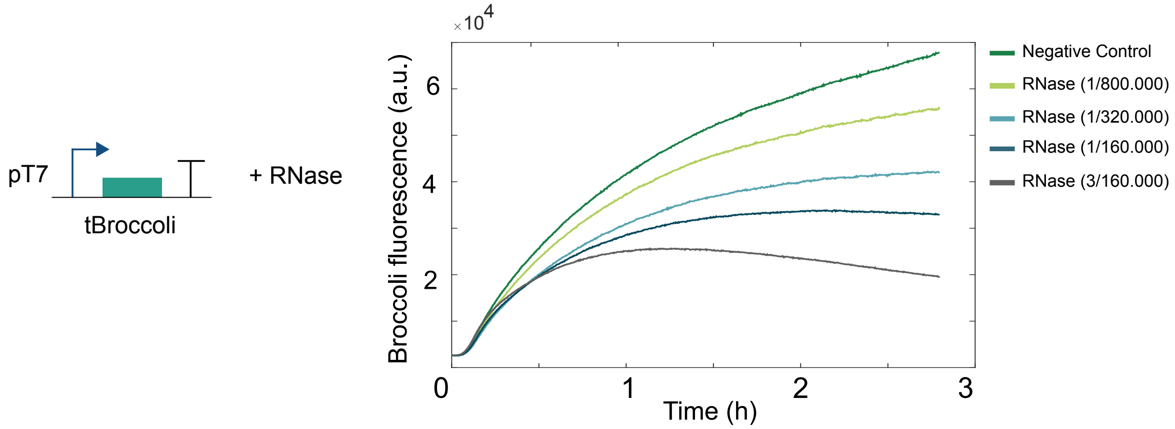


Supplementary Figure 7: Characterization of RNase molecules using tBroccoli aptamer. Different concentrations of RNAse molecules were tested to obtain a balance between production and degradation of RNA molecules. Negative control represents the fluorescent aptamer without RNAse. Dilutions were calculated based on the commercial RNAse (Methods).

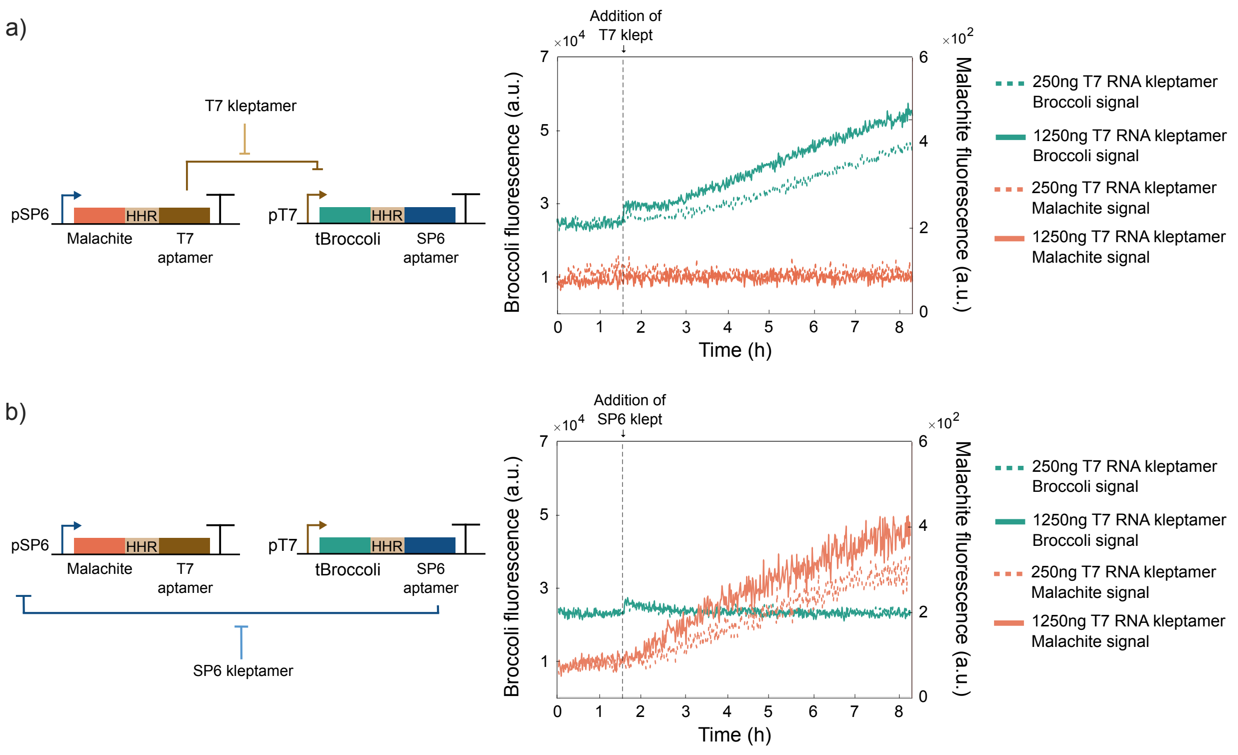


Supplementary Figure 8: Performance of trans-acting circuits using RNA kleptamers. **a)** State A of the RNA-based toggle switch where the expression of T7 inhibitory aptamer is repressing tBroccoli. The addition of T7 RNA kleptamer at 250ng and 1250ng increased tBroccoli fluorescence signal. **b)** State B of the RNA-based toggle switch where the expression of SP6 inhibitory aptamer is repressing Malachite Green aptamer. The addition of SP6 RNA kleptamer at 250ng and 1250ng increased Malachite Green fluorescence signal. Spikes in fluorescence are due to injection-induced temperature decrease.
